# Severe bottleneck of ancient *Homo* populations: Insights from computational modeling and fossil evidence

**DOI:** 10.1101/2025.02.01.636025

**Authors:** Sumei Zhou, Yiwen Zhou, Fabio Di Vincenzo, Giorgio Manzi, Wangjie Hu, Ziqian Hao, Zhen Shao, Yi-Hsuan Pan, Haipeng Li

**Author notes:** Corresponding authors: Yi-Hsuan Pan, and Haipeng Li, **Email:**. These authors contributed equally to this work (Sumei Zhou, Yiwen Zhou). **Author Contributions:** Conceptualization: SZ, YZ, FDV, GM, WH, ZH, YHP, HL; Investigation: SZ, YZ, FDV; Supervision: GM, YHP, HL; Writing: SZ, YZ, FDV, GM, YHP, HL. **Competing Interest Statement:** Authors declare that they have no competing interests.

## Abstract

Reconstructing ancient population size history is essential for understanding the evolutionary origin of *Homo sapiens*. Using the fast infinitesimal time coalescent process (FitCoal), which precisely computes the expected site frequency spectrum (SFS), we revisited the debated population bottleneck that occurred approximately 930 thousand years ago. By benchmarking FitCoal against *mushi* and ten billion coalescent simulations, we demonstrate that FitCoal achieves both superior speed and accuracy in expected SFS estimation. Analyses of simulated datasets confirmed that FitCoal reliably recovered the bottleneck, whereas *mushi* failed under identical conditions. Independent fossil and paleoclimate evidence further corroborates the timing and evolutionary impact of this bottleneck, linking it to hominin dispersals, speciation events, and a subsequent increase in brain size. These findings refine the demographic history of *Homo* during the Pleistocene and highlight the importance of high-precision SFS computation for revealing critical evolutionary transitions that shaped modern human ancestry.

## Introduction

Over the past four million years, at least 22 hominin taxa have emerged, yet only modern humans (*Homo sapiens*) have persisted to the present day. The Early and Middle Pleistocene epochs represent a pivotal period in *Homo* evolution, during which African *H. erectus* declined (Di Vincenzo and Manzi 2023) and gave rise to a new species ancestral to modern humans, Neanderthals, and Denisovans (Prüfer, et al. 2014). Understanding the evolutionary trajectory leading to modern humans therefore requires reconstructing population size dynamics throughout the Pleistocene.

Site frequency spectrum (SFS)-based methods provide powerful tools for reconstructing population histories (Li and Stephan 2006; DeWitt, et al. 2021; Excoffier, et al. 2021). The SFS comprises (*n*–1) allele frequency categories, where *n* is the number of sampled sequences. Previous studies developed methods to estimate the expected branch lengths for each SFS category and calculate the likelihoods of observed SFS data under various demographic scenarios. However, these methods may lack sufficient accuracy for inferring the detailed population history of *Homo* during the Pleistocene. To overcome this limitation, the fast infinitesimal time coalescent process (FitCoal) was recently introduced to precisely infer both recent and ancient population size histories using the SFS (Hu, et al. 2023). FitCoal accurately calculates the expected SFS and associated likelihood. As different theoretical demographic models may yield similar SFS patterns (Myers, et al. 2008), FitCoal prioritizes biologically plausible scenarios and systematically increases model complexity to identify the most appropriate demographic model. Moreover, high-frequency alleles within the SFS are particularly susceptible to confounding factors, such as positive selection (Li 2010; Vy and Kim 2015), introgression (Tagore and Akey 2025), population structure (Kern and Hey 2017), and misclassification of ancestral and derived variants (Baudry and Depaulis 2003). Consequently, excluding such overrepresented alleles reduces biases and enhances the robustness and reliability of inferred demographic histories (Bhaskar, et al. 2015; Liu and Fu 2015; Hu, et al. 2023).

## Results and Discussion

It has been widely recognized for nearly four decades that current African and non-African human populations share a common evolutionary history prior to their divergence. Our previous research identified a severe population bottleneck occurring approximately 930 thousand years ago (Ka) in both 12 African and 19 non-African populations (Hu, et al. 2023). However, recent studies have challenged this finding, suggesting that this bottleneck was confined to African populations alone and absent in non-African populations (Cousins and Durvasula 2025; Deng, et al. 2025). To resolve this discrepancy, we re-examined our earlier analyses and confirmed that both the timing and severity of the bottleneck were virtually identical between African and non-African populations (Fig. 1A, B).

**Fig 1.**
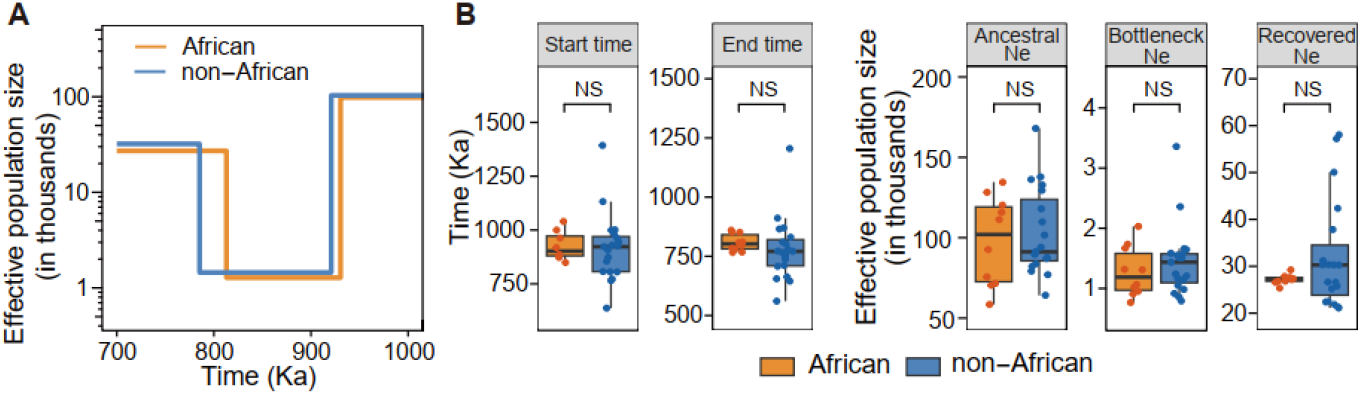
The ancient human severe bottleneck. (**A**) The severe bottleneck inferred in African and non-African populations. Mean values are shown. (**B**) Estimated bottleneck parameters for 10 African and 19 non-African populations. Two South African populations (San and Bantu) were excluded due to small sample sizes. Estimates are based on previously published results (Hu, et al. 2023). Two-tailed Student’s *t*-test; NS, not significant.

To further validate the efficacy of FitCoal, we compared inferred population histories obtained using both the complete site frequency spectrum (SFS) and a restricted subset of the SFS under an arbitrary bottleneck model. Remarkably, the bottleneck was accurately inferred even when analyzing only 10% of the SFS categories (minor allele frequencies ≤0.1) (Fig. 2A). FitCoal effectively captures subtle yet widespread signals present in the SFS due to the bottleneck event, as each coalescent state maintains a non-zero probability when traced back to this event (Hu, et al. 2023). This attribute significantly reduces biases from confounding factors, which predominantly affect high-frequency alleles. Although recent work partially addressed this issue by simulating ancestral and derived variant misclassification, the approach inadvertently increased model complexity (Cousins and Durvasula 2025). In contrast, FitCoal mitigates these confounding influences by specifically excluding high-frequency alleles (Bhaskar, et al. 2015; Liu and Fu 2015). These results highlight the abundant demographic information contained within the SFS and emphasize the robustness of FitCoal in accurately reconstructing complex evolutionary histories.

**Fig. 2.**
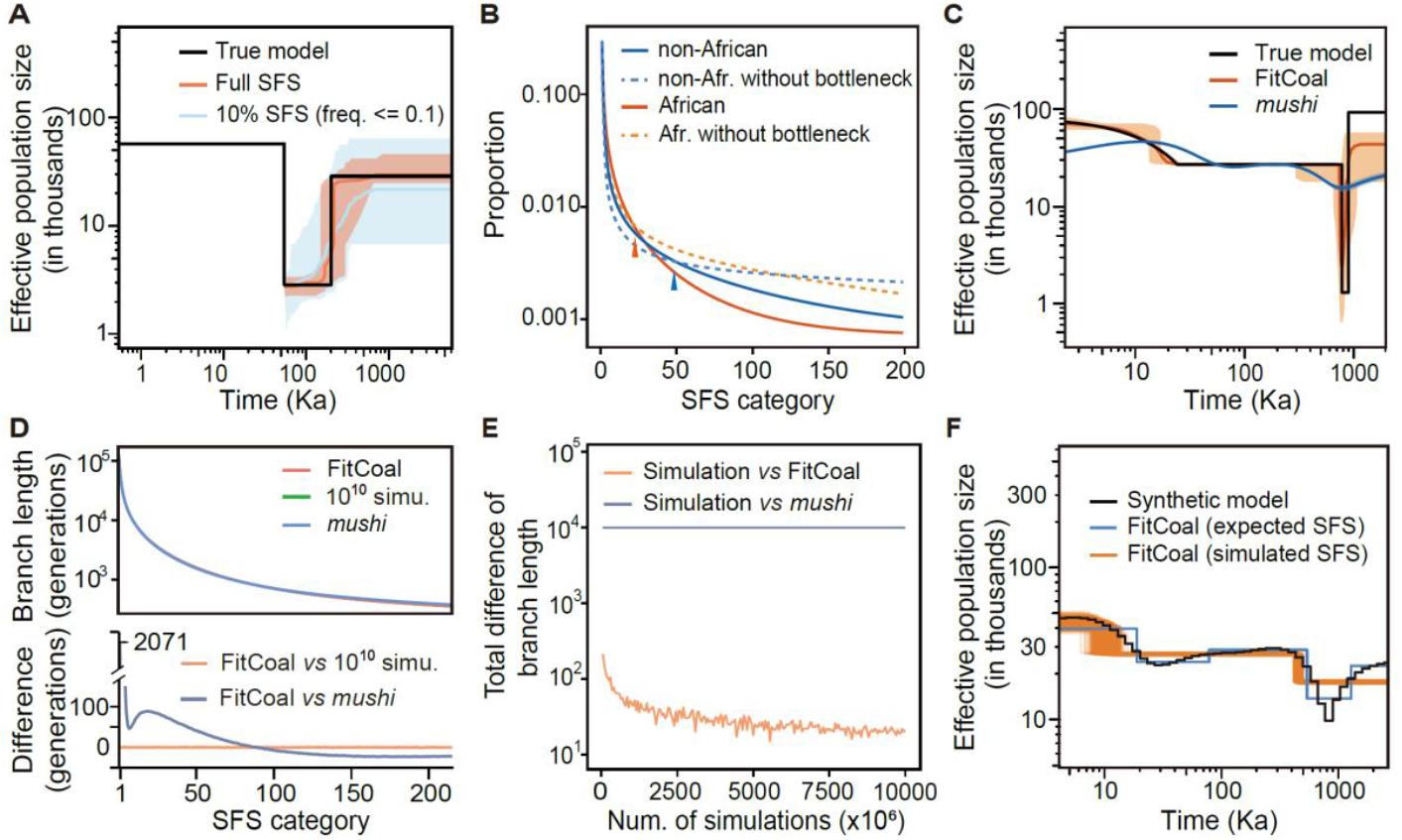
Implications of FitCoal’s high-precision computation. (**A**) FitCoal-inferred population size histories using complete-SFS and minor alleles (*freq* ≤0.1). Colored bands represent estimated ranges spanning the 2.5th to the 97.5th percentiles. (**B)** Impact of the ancient severe bottleneck on the SFS. Solid orange and blue lines show the normalized expected SFS under African and non-African the demographic histories, respectively, while dashed lines represent cases without the ancient severe bottleneck. Arrows indicate the onset of allele deficiency. (**C**) Population size histories inferred by FitCoal and *mushi*. Shaded areas denote estimated ranges between the 2.5th and 97.5th percentiles. The black line represents the true model, mimicking the demographic history of African agriculturalist populations. (**D**) Upper panel: expected branch lengths for each SFS category calculated by FitCoal, *mushi*, and 1 × 10^10^ *msprime* simulations under the synthetic model. Lower panel: deviations from FitCoal estimates. (**E**) Effect of simulation number on the accuracy of averaged branch lengths, compared with FitCoal and *mushi* estimates under the synthetic model. (**F**) FitCoal-inferred population size histories from simulated and the expected SFSs under the synthetic model. Red lines indicate histories inferred from 1,000 simulated datasets (*n*=200, *L*=100 Mb), and the blue line represents the history inferred from the expected SFS.

We further investigated the changes in the SFS induced by the presence or absence of the ancient severe bottleneck event. This bottleneck significantly reduced the proportions of mid- and high-frequency alleles in both African and non-African populations (Fig. 2B). Notably, the reduction was more pronounced in African populations, becoming evident from the 20th SFS category onward, compared to the 50th category in non-African populations. This finding is consistent with the broader range of bottleneck estimates observed in non-African populations (Fig. 1B). To further evaluate the detectability of the bottleneck, we simulated genome-wide polymorphic datasets based on inferred demographic histories from African populations. FitCoal successfully recovered the bottleneck signal (Fig. 2C), whereas *mushi* (DeWitt, et al. 2021) did not detect this signal under the same conditions. The log-likelihood of the demographic history inferred by FitCoal was also significantly higher than that inferred by *mushi* (-1211.63 *vs* - 2849.65, *P*-value < 2.2× 10^−16^, Fig. 3). These outcomes reinforce FitCoal’s effectiveness and reliability in identifying the ancient severe bottleneck from genomic data.

**Fig. 3.**
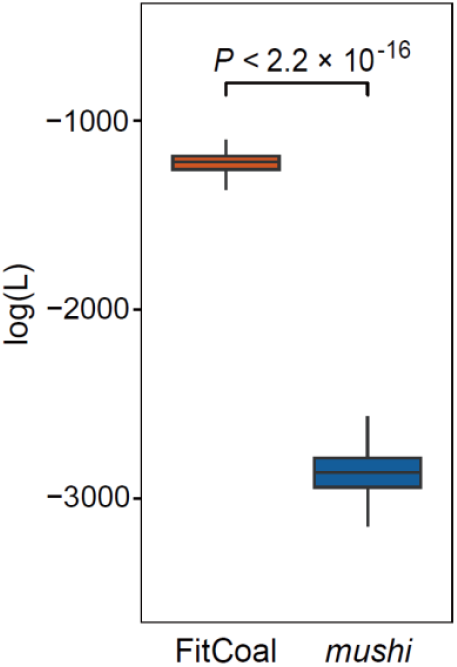
Log-likelihood values of FitCoal- and *mushi*-inferred histories. The true model is shown in Fig. 2C. Demographic histories were inferred from 1,000 simulated genome-wide datasets using FitCoal and *mushi*. Statistical significance was determined by two-tailed Student’s *t*-test.

To further explore the differences between FitCoal and *mushi* in inferring population history, we compared their estimations of expected branch lengths, a critical component in calculating the expected SFS. It is known that *mushi* relies on the joint probability density function of coalescent times, making it susceptible to accumulated numerical errors (Polanski, et al. 2003; Polanski and Kimmel 2003; Živković and Wiehe 2008). FitCoal overcomes this limitation by introducing a novel framework, the fast infinitesimal time coalescent process. For a fair comparison, we examined a synthetic demographic model previously associated with misidentification of a severe bottleneck (Deng, et al. 2025). The expected branch lengths estimated by FitCoal and *mushi* differed by up to 6.21%, corresponding to a cumulative discrepancy of 9,981 generations across SFS categories (Fig. 2D). To asses which method more closely reflects true values, we conducted 1 × 10^10^ *msprime* coalescent simulations (Baumdicker, et al. 2022) as a benchmark. FitCoal’s estimates matched simulated values exactly in 194 of 215 SFS categories (90.2%), with only a single-generation deviation in the remaining categories (Table 1). Conversely, *mushi* exhibited deviations two orders of magnitude larger (Fig. 2E), underscoring FitCoal’s superior accuracy. A recent study reported no apparent differences between *mushi* and *msprime* simulations (Cousins and Durvasula 2025), likely because their analyses were limited to comparisons analogous to the upper panel of Figure 2D, where differences remain minimal. Furthermore, FitCoal demonstrated remarkable computational efficiency, completing the calculation in only 0.00013 seconds, whereas *mushi* and *msprime* required 1.29 seconds and 4.90 years, respectively. These results highlight FitCoal’s advantages in both accuracy and computational speed.

**Table 1.** Expected branch lengths for each SFS category.

| Type<sup>1</sup> | FitCoal<br>(Generations) | $1 \times 10^{10}$<br>simulations<br>(Generations) | *Mushi*<br>(Generations) | Absolute diff.<br>FitCoal vs<br>simulations | Absolute diff.<br>FitCoal vs<br>*mushi* |
| --- | --- | --- | --- | --- | --- |
| 1 | 129059 | 129059 | 126988 | 0 (0.00%) | 2071 (1.60%) |
| 2 | 57514 | 57514 | 56423 | 0 (0.00%) | 1091 (1.90%) |
| 3 | 35743 | 35743 | 35270 | 0 (0.00%) | 473 (1.32%) |
| 4 | 25931 | 25931 | 25730 | 0 (0.00%) | 201 (0.78%) |
| 5 | 20441 | 20441 | 20349 | 0 (0.00%) | 92 (0.45%) |
| 6 | 16922 | 16922 | 16867 | 0 (0.00%) | 55 (0.33%) |
| 7 | 14455 | 14455 | 14409 | 0 (0.00%) | 46 (0.32%) |
| 8 | 12617 | 12617 | 12568 | 0 (0.00%) | 49 (0.39%) |
| 9 | 11187 | 11187 | 11132 | 0 (0.00%) | 55 (0.49%) |
| 10 | 10039 | 10039 | 9977 | 0 (0.00%) | 62 (0.62%) |
| 11 | 9096 | 9096 | 9026 | 0 (0.00%) | 70 (0.77%) |
| 12 | 8306 | 8306 | 8231 | 0 (0.00%) | 75 (0.90%) |
| 13 | 7634 | 7634 | 7554 | 0 (0.00%) | 80 (1.05%) |
| 14 | 7056 | 7056 | 6972 | 0 (0.00%) | 84 (1.19%) |
| 15 | 6552 | 6552 | 6466 | 0 (0.00%) | 86 (1.31%) |
| 16 | 6110 | 6110 | 6023 | 0 (0.00%) | 87 (1.42%) |
| 17 | 5719 | 5719 | 5631 | 0 (0.00%) | 88 (1.54%) |
| 18 | 5371 | 5371 | 5282 | 0 (0.00%) | 89 (1.66%) |
| 19 | 5059 | 5059 | 4970 | 0 (0.00%) | 89 (1.76%) |
| 20 | 4778 | 4778 | 4689 | 0 (0.00%) | 89 (1.86%) |
| 21 | 4523 | 4523 | 4435 | 0 (0.00%) | 88 (1.95%) |
| 22 | 4291 | 4291 | 4204 | 0 (0.00%) | 87 (2.03%) |
| 23 | 4080 | 4080 | 3994 | 0 (0.00%) | 86 (2.11%) |
| 24 | 3886 | 3886 | 3801 | 0 (0.00%) | 85 (2.19%) |
| 25 | 3708 | 3708 | 3624 | 0 (0.00%) | 84 (2.27%) |
| 26 | 3544 | 3543 | 3461 | 1 (0.03%) | 83 (2.34%) |
| 27 | 3392 | 3392 | 3311 | 0 (0.00%) | 81 (2.39%) |
| 28 | 3251 | 3251 | 3172 | 0 (0.00%) | 79 (2.43%) |
| 29 | 3120 | 3120 | 3042 | 0 (0.00%) | 78 (2.50%) |
| 30 | 2998 | 2998 | 2922 | 0 (0.00%) | 76 (2.54%) |
| 31 | 2884 | 2884 | 2810 | 0 (0.00%) | 74 (2.57%) |
| 32 | 2777 | 2777 | 2705 | 0 (0.00%) | 72 (2.59%) |
| 33 | 2678 | 2677 | 2607 | 1 (0.04%) | 71 (2.65%) |
| 34 | 2584 | 2584 | 2515 | 0 (0.00%) | 69 (2.67%) |
| 35 | 2496 | 2496 | 2429 | 0 (0.00%) | 67 (2.68%) |
| 36 | 2413 | 2413 | 2348 | 0 (0.00%) | 65 (2.69%) |
| 37 | 2334 | 2334 | 2271 | 0 (0.00%) | 63 (2.70%) |
| 38 | 2260 | 2260 | 2199 | 0 (0.00%) | 61 (2.70%) |
| 39 | 2190 | 2190 | 2131 | 0 (0.00%) | 59 (2.69%) |
| 40 | 2124 | 2124 | 2066 | 0 (0.00%) | 58 (2.73%) |
| 41 | 2062 | 2061 | 2005 | 1 (0.05%) | 57 (2.76%) |
| 42 | 2002 | 2002 | 1947 | 0 (0.00%) | 55 (2.75%) |
| 43 | 1945 | 1945 | 1892 | 0 (0.00%) | 53 (2.72%) |
| 44 | 1892 | 1891 | 1840 | 1 (0.05%) | 52 (2.75%) |
| 45 | 1840 | 1840 | 1791 | 0 (0.00%) | 49 (2.66%) |
| 46 | 1791 | 1792 | 1743 | 1 (0.06%) | 48 (2.68%) |
| 47 | 1745 | 1745 | 1698 | 0 (0.00%) | 47 (2.69%) |
| 48 | 1700 | 1701 | 1656 | 1 (0.06%) | 44 (2.59%) |
| 49 | 1658 | 1658 | 1615 | 0 (0.00%) | 43 (2.59%) |
| 50 | 1617 | 1617 | 1576 | 0 (0.00%) | 41 (2.54%) |
| 51 | 1579 | 1578 | 1538 | 1 (0.06%) | 41 (2.60%) |
| 52 | 1541 | 1541 | 1503 | 0 (0.00%) | 38 (2.47%) |
| 53 | 1506 | 1506 | 1468 | 0 (0.00%) | 38 (2.52%) |
| 54 | 1472 | 1471 | 1436 | 1 (0.07%) | 36 (2.45%) |
| 55 | 1439 | 1439 | 1404 | 0 (0.00%) | 35 (2.43%) |
| 56 | 1407 | 1407 | 1374 | 0 (0.00%) | 33 (2.35%) |
| 57 | 1377 | 1377 | 1345 | 0 (0.00%) | 32 (2.32%) |
| 58 | 1348 | 1348 | 1318 | 0 (0.00%) | 30 (2.23%) |
| 59 | 1320 | 1320 | 1291 | 0 (0.00%) | 29 (2.20%) |
| 60 | 1293 | 1293 | 1265 | 0 (0.00%) | 28 (2.17%) |
| 61 | 1267 | 1267 | 1241 | 0 (0.00%) | 26 (2.05%) |
| 62 | 1242 | 1242 | 1217 | 0 (0.00%) | 25 (2.01%) |
| 63 | 1218 | 1218 | 1194 | 0 (0.00%) | 24 (1.97%) |
| 64 | 1195 | 1195 | 1172 | 0 (0.00%) | 23 (1.92%) |
| 65 | 1173 | 1173 | 1151 | 0 (0.00%) | 22 (1.88%) |
| 66 | 1151 | 1151 | 1131 | 0 (0.00%) | 20 (1.74%) |
| 67 | 1131 | 1131 | 1111 | 0 (0.00%) | 20 (1.77%) |
| 68 | 1110 | 1111 | 1092 | 1 (0.09%) | 18 (1.62%) |
| 69 | 1091 | 1091 | 1074 | 0 (0.00%) | 17 (1.56%) |
| 70 | 1072 | 1072 | 1056 | 0 (0.00%) | 16 (1.49%) |
| 71 | 1054 | 1054 | 1039 | 0 (0.00%) | 15 (1.42%) |
| 72 | 1036 | 1037 | 1022 | 1 (0.10%) | 14 (1.35%) |
| 73 | 1019 | 1020 | 1006 | 1 (0.10%) | 13 (1.28%) |
| 74 | 1003 | 1003 | 991 | 0 (0.00%) | 12 (1.20%) |
| 75 | 987 | 987 | 976 | 0 (0.00%) | 11 (1.11%) |
| 76 | 972 | 972 | 961 | 0 (0.00%) | 11 (1.13%) |
| 77 | 957 | 957 | 947 | 0 (0.00%) | 10 (1.04%) |
| 78 | 942 | 942 | 934 | 0 (0.00%) | 8 (0.85%) |
| 79 | 928 | 928 | 921 | 0 (0.00%) | 7 (0.75%) |
| 80 | 914 | 914 | 908 | 0 (0.00%) | 6 (0.66%) |
| 81 | 901 | 901 | 895 | 0 (0.00%) | 6 (0.67%) |
| 82 | 888 | 888 | 883 | 0 (0.00%) | 5 (0.56%) |
| 83 | 876 | 876 | 872 | 0 (0.00%) | 4 (0.46%) |
| 84 | 864 | 864 | 860 | 0 (0.00%) | 4 (0.46%) |
| 85 | 852 | 852 | 849 | 0 (0.00%) | 3 (0.35%) |
| 86 | 840 | 840 | 839 | 0 (0.00%) | 1 (0.12%) |
| 87 | 829 | 829 | 828 | 0 (0.00%) | 1 (0.12%) |
| 88 | 819 | 819 | 818 | 0 (0.00%) | 1 (0.12%) |
| 89 | 808 | 808 | 808 | 0 (0.00%) | 0 (0.00%) |
| 90 | 798 | 798 | 799 | 0 (0.00%) | 1 (0.13%) |
| 91 | 788 | 788 | 789 | 0 (0.00%) | 1 (0.13%) |
| 92 | 778 | 778 | 780 | 0 (0.00%) | 2 (0.26%) |
| 93 | 769 | 769 | 771 | 0 (0.00%) | 2 (0.26%) |
| 94 | 759 | 759 | 763 | 0 (0.00%) | 4 (0.53%) |
| 95 | 751 | 750 | 755 | 1 (0.13%) | 4 (0.53%) |
| 96 | 742 | 742 | 746 | 0 (0.00%) | 4 (0.54%) |
| 97 | 733 | 733 | 738 | 0 (0.00%) | 5 (0.68%) |
| 98 | 725 | 725 | 731 | 0 (0.00%) | 6 (0.83%) |
| 99 | 717 | 717 | 723 | 0 (0.00%) | 6 (0.84%) |
| 100 | 709 | 709 | 716 | 0 (0.00%) | 7 (0.99%) |
| 101 | 701 | 701 | 709 | 0 (0.00%) | 8 (1.14%) |
| 102 | 694 | 694 | 702 | 0 (0.00%) | 8 (1.15%) |
| 103 | 686 | 686 | 695 | 0 (0.00%) | 9 (1.31%) |
| 104 | 679 | 680 | 688 | 1 (0.15%) | 9 (1.33%) |
| 105 | 672 | 672 | 682 | 0 (0.00%) | 10 (1.49%) |
| 106 | 666 | 665 | 675 | 1 (0.15%) | 9 (1.35%) |
| 107 | 659 | 659 | 669 | 0 (0.00%) | 10 (1.52%) |
| 108 | 652 | 653 | 663 | 1 (0.15%) | 11 (1.69%) |
| 109 | 646 | 646 | 657 | 0 (0.00%) | 11 (1.70%) |
| 110 | 640 | 640 | 651 | 0 (0.00%) | 11 (1.72%) |
| 111 | 634 | 634 | 646 | 0 (0.00%) | 12 (1.89%) |
| 112 | 628 | 628 | 640 | 0 (0.00%) | 12 (1.91%) |
| 113 | 622 | 622 | 635 | 0 (0.00%) | 13 (2.09%) |
| 114 | 616 | 616 | 629 | 0 (0.00%) | 13 (2.11%) |
| 115 | 611 | 611 | 624 | 0 (0.00%) | 13 (2.13%) |
| 116 | 605 | 605 | 619 | 0 (0.00%) | 14 (2.31%) |
| 117 | 600 | 600 | 614 | 0 (0.00%) | 14 (2.33%) |
| 118 | 595 | 595 | 609 | 0 (0.00%) | 14 (2.35%) |
| 119 | 590 | 590 | 605 | 0 (0.00%) | 15 (2.54%) |
| 120 | 585 | 585 | 600 | 0 (0.00%) | 15 (2.56%) |
| 121 | 580 | 580 | 595 | 0 (0.00%) | 15 (2.59%) |
| 122 | 575 | 575 | 591 | 0 (0.00%) | 16 (2.78%) |
| 123 | 571 | 570 | 587 | 1 (0.18%) | 16 (2.80%) |
| 124 | 566 | 566 | 582 | 0 (0.00%) | 16 (2.83%) |
| 125 | 561 | 561 | 578 | 0 (0.00%) | 17 (3.03%) |
| 126 | 557 | 557 | 574 | 0 (0.00%) | 17 (3.05%) |
| 127 | 553 | 553 | 570 | 0 (0.00%) | 17 (3.07%) |
| 128 | 549 | 549 | 566 | 0 (0.00%) | 17 (3.10%) |
| 129 | 544 | 544 | 562 | 0 (0.00%) | 18 (3.31%) |
| 130 | 540 | 540 | 558 | 0 (0.00%) | 18 (3.33%) |
| 131 | 536 | 536 | 554 | 0 (0.00%) | 18 (3.36%) |
| 132 | 532 | 532 | 551 | 0 (0.00%) | 19 (3.57%) |
| 133 | 529 | 529 | 547 | 0 (0.00%) | 18 (3.40%) |
| 134 | 525 | 525 | 544 | 0 (0.00%) | 19 (3.62%) |
| 135 | 521 | 521 | 540 | 0 (0.00%) | 19 (3.65%) |
| 136 | 517 | 517 | 537 | 0 (0.00%) | 20 (3.87%) |
| 137 | 514 | 514 | 533 | 0 (0.00%) | 19 (3.70%) |
| 138 | 510 | 510 | 530 | 0 (0.00%) | 20 (3.92%) |
| 139 | 507 | 507 | 527 | 0 (0.00%) | 20 (3.94%) |
| 140 | 504 | 504 | 523 | 0 (0.00%) | 19 (3.77%) |
| 141 | 500 | 500 | 520 | 0 (0.00%) | 20 (4.00%) |
| 142 | 497 | 497 | 517 | 0 (0.00%) | 20 (4.02%) |
| 143 | 494 | 494 | 514 | 0 (0.00%) | 20 (4.05%) |
| 144 | 491 | 491 | 511 | 0 (0.00%) | 20 (4.07%) |
| 145 | 488 | 488 | 508 | 0 (0.00%) | 20 (4.10%) |
| 146 | 485 | 485 | 505 | 0 (0.00%) | 20 (4.12%) |
| 147 | 482 | 482 | 503 | 0 (0.00%) | 21 (4.36%) |
| 148 | 479 | 479 | 500 | 0 (0.00%) | 21 (4.38%) |
| 149 | 476 | 476 | 497 | 0 (0.00%) | 21 (4.41%) |
| 150 | 473 | 473 | 494 | 0 (0.00%) | 21 (4.44%) |
| 151 | 470 | 470 | 492 | 0 (0.00%) | 22 (4.68%) |
| 152 | 467 | 467 | 489 | 0 (0.00%) | 22 (4.71%) |
| 153 | 465 | 465 | 486 | 0 (0.00%) | 21 (4.52%) |
| 154 | 462 | 462 | 484 | 0 (0.00%) | 22 (4.76%) |
| 155 | 460 | 459 | 481 | 1 (0.22%) | 21 (4.57%) |
| 156 | 457 | 457 | 479 | 0 (0.00%) | 22 (4.81%) |
| 157 | 454 | 454 | 476 | 0 (0.00%) | 22 (4.85%) |
| 158 | 452 | 452 | 474 | 0 (0.00%) | 22 (4.87%) |
| 159 | 449 | 449 | 472 | 0 (0.00%) | 23 (5.12%) |
| 160 | 447 | 447 | 469 | 0 (0.00%) | 22 (4.92%) |
| 161 | 445 | 445 | 467 | 0 (0.00%) | 22 (4.94%) |
| 162 | 442 | 442 | 465 | 0 (0.00%) | 23 (5.20%) |
| 163 | 440 | 440 | 462 | 0 (0.00%) | 22 (5.00%) |
| 164 | 438 | 438 | 460 | 0 (0.00%) | 22 (5.02%) |
| 165 | 436 | 435 | 458 | 1 (0.23%) | 22 (5.05%) |
| 166 | 433 | 433 | 456 | 0 (0.00%) | 23 (5.31%) |
| 167 | 431 | 431 | 454 | 0 (0.00%) | 23 (5.34%) |
| 168 | 429 | 429 | 452 | 0 (0.00%) | 23 (5.36%) |
| 169 | 427 | 427 | 450 | 0 (0.00%) | 23 (5.39%) |
| 170 | 425 | 425 | 447 | 0 (0.00%) | 22 (5.18%) |
| 171 | 423 | 423 | 445 | 0 (0.00%) | 22 (5.20%) |
| 172 | 421 | 421 | 443 | 0 (0.00%) | 22 (5.23%) |
| 173 | 419 | 419 | 441 | 0 (0.00%) | 22 (5.25%) |
| 174 | 417 | 417 | 440 | 0 (0.00%) | 23 (5.52%) |
| 175 | 415 | 415 | 438 | 0 (0.00%) | 23 (5.54%) |
| 176 | 413 | 413 | 436 | 0 (0.00%) | 23 (5.57%) |
| 177 | 411 | 411 | 434 | 0 (0.00%) | 23 (5.60%) |
| 178 | 409 | 409 | 432 | 0 (0.00%) | 23 (5.62%) |
| 179 | 407 | 407 | 430 | 0 (0.00%) | 23 (5.65%) |
| 180 | 406 | 406 | 428 | 0 (0.00%) | 22 (5.42%) |
| 181 | 404 | 404 | 427 | 0 (0.00%) | 23 (5.69%) |
| 182 | 402 | 402 | 425 | 0 (0.00%) | 23 (5.72%) |
| 183 | 400 | 400 | 423 | 0 (0.00%) | 23 (5.75%) |
| 184 | 399 | 398 | 421 | 1 (0.25%) | 22 (5.51%) |
| 185 | 397 | 397 | 420 | 0 (0.00%) | 23 (5.79%) |
| 186 | 395 | 395 | 418 | 0 (0.00%) | 23 (5.82%) |
| 187 | 393 | 393 | 416 | 0 (0.00%) | 23 (5.85%) |
| 188 | 392 | 392 | 415 | 0 (0.00%) | 23 (5.87%) |
| 189 | 390 | 390 | 413 | 0 (0.00%) | 23 (5.90%) |
| 190 | 389 | 389 | 411 | 0 (0.00%) | 22 (5.66%) |
| 191 | 387 | 387 | 410 | 0 (0.00%) | 23 (5.94%) |
| 192 | 386 | 386 | 408 | 0 (0.00%) | 22 (5.70%) |
| 193 | 384 | 384 | 407 | 0 (0.00%) | 23 (5.99%) |
| 194 | 382 | 382 | 405 | 0 (0.00%) | 23 (6.02%) |
| 195 | 381 | 381 | 404 | 0 (0.00%) | 23 (6.04%) |
| 196 | 379 | 379 | 402 | 0 (0.00%) | 23 (6.07%) |
| 197 | 378 | 378 | 401 | 0 (0.00%) | 23 (6.08%) |
| 198 | 377 | 377 | 399 | 0 (0.00%) | 22 (5.84%) |
| 199 | 375 | 375 | 398 | 0 (0.00%) | 23 (6.13%) |
| 200 | 374 | 374 | 396 | 0 (0.00%) | 22 (5.88%) |
| 201 | 372 | 372 | 395 | 0 (0.00%) | 23 (6.18%) |
| 202 | 371 | 371 | 393 | 0 (0.00%) | 22 (5.93%) |
| 203 | 370 | 369 | 392 | 1 (0.27%) | 22 (5.95%) |
| 204 | 368 | 368 | 390 | 0 (0.00%) | 22 (5.98%) |
| 205 | 367 | 367 | 389 | 0 (0.00%) | 22 (5.99%) |
| 206 | 365 | 365 | 388 | 0 (0.00%) | 23 (6.30%) |
| 207 | 364 | 364 | 386 | 0 (0.00%) | 22 (6.04%) |
| 208 | 363 | 363 | 385 | 0 (0.00%) | 22 (6.06%) |
| 209 | 362 | 361 | 384 | 1 (0.28%) | 22 (6.08%) |
| 210 | 360 | 360 | 382 | 0 (0.00%) | 22 (6.11%) |
| 211 | 359 | 359 | 381 | 0 (0.00%) | 22 (6.13%) |
| 212 | 358 | 358 | 380 | 0 (0.00%) | 22 (6.15%) |
| 213 | 357 | 357 | 378 | 0 (0.00%) | 21 (5.88%) |
| 214 | 355 | 355 | 377 | 0 (0.00%) | 22 (6.20%) |
| 215 | 354 | 354 | 376 | 0 (0.00%) | 22 (6.21%) |
| Total diff. | – | – | – | 21 | 9,981 |

<sup>1</sup>The number of sampled sequences is 216, following Deng *et al* (2025).

The observed, not the expected, SFSs were used as input for FitCoal (Hu, et al. 2023). A recent study input an expected SFS, leading to the detection of a severe-bottleneck-like event (Deng, et al. 2025). To clarify this issue, we compared expected and observed SFS and found that reliably estimating expected branch lengths requires at least 1 × 10^10^ simulations (Fig. 2E) and that the genome-wide observed SFS varies significantly among population samples. Considering this inherent variability, FitCoal employs a conservative default threshold of 20% likelihood gain, minimizing false positives in demographic model selection (Hu, et al. 2023). We validated this conservativity by simulating 1,000 genome-wide polymorphic datasets under the previously studied synthetic model (Deng, et al. 2025). FitCoal consistently avoided overfitting the mild population decrease (Fig. 2F), as the conservative threshold effectively minimized false positives. When using the expected SFS as input, a lower threshold (1% likelihood gain) was required to achieve optimized likelihoods without premature termination, yielding accurate demographic histories (Fig. 2F). These findings show the importance of clearly distinguishing between expected and observed SFSs in demographic analyses.

Existence of the ancient severe bottleneck is supported by multiple lines of evidence, including gaps in the African hominin fossil record (Profico, et al. 2016), the timing of ancestral chromosome fusion events (Dreszer, et al. 2007; Poszewiecka, et al. 2022; Poszewiecka, et al. 2023), major cultural and behavioral shifts (Clark and Linares-Matás 2024), climatic transitions during the Early-to-Middle Pleistocene (Jouzel, et al. 2007), and fluctuations in mammalian speciation and extinction rates (Cohen, et al. 2022). Moreover, the timing of the bottleneck coincides with major episodes of hominin dispersals from Africa to Eurasia (Muttoni and Kent 2024), and the evolutionary context inferred from the earliest known human facial fossil in Western Europe (Huguet, et al. 2025) aligns closely with the demographic decline scenario associated with this bottleneck.

Between 1,000 and 100 Ka, at least two significant macroevolutionary events occurred (Fig. 4A), coinciding with the “muddle in the middle”, a period characterized by hominin coexistence and morphological diversity (Bergström, et al. 2021; Ni, et al. 2021; Fu, et al. 2025). Since small population sizes facilitate speciation (Gould and Eldredge 1977), our findings imply that this severe bottleneck significantly impacted phenotypic and behavioral evolution in ancestral humans. Particularly, its role in adaptation and speciation warrant further investigation. The fixation of chromosome 2 fusion and its possible contribution to reproductive isolation remains unresolved. To further explore this, we analyzed endocranial volumes and encephalization quotient data from African fossil specimens. Our analysis revealed a significant increase in brain size following the bottleneck (Fig. 4B, Table 2), suggesting a potential relationship between population contraction and accelerated cognitive and social evolution. Additionally, recent evidence indicates that the last populations of African *H. erectus* experienced extreme aridity at the onset of the bottleneck, potentially driving behavioral adaptations critical for improved survival and resource utilization (Mercader, et al. 2025).

**Table 2.** Endocranial volume and the encephalization quotient of African *Homo erectus* and the LCA.

| Taxon | Specimen | Endocranial volume (cm<sup>3</sup>) | Encephalization quotient |
| --- | --- | --- | --- |
| *H. erectus*<br>(Africa) | KNM-ER 3732 | 750 | 4.83 |
|  | KNM-ER 3733 | 848 | 5.46 |
|  | KNM-ER 3883 | 804 | 5.18 |
|  | Daka | 995 | 6.41 |
|  | Buia | 995 | 6.41 |
|  | KNM-ER 42700 | 721 | 4.64 |
|  | KNM-WT 15000 | 900 | 5.79 |
|  | OH 9 | 1,067 | 6.87 |
|  | OH 12 | 727 | 4.68 |
| LCA | Bodo | 1,250 | 6.80 |
|  | Broken Hill 1 | 1,325 | 7.21 |
|  | Saldanha | 1,225 | 6.66 |
|  | Omo-Kibish 2 | 1,430 | 7.77 |
|  | Ndutu | 1,100 | 5.98 |

**Fig. 4.**
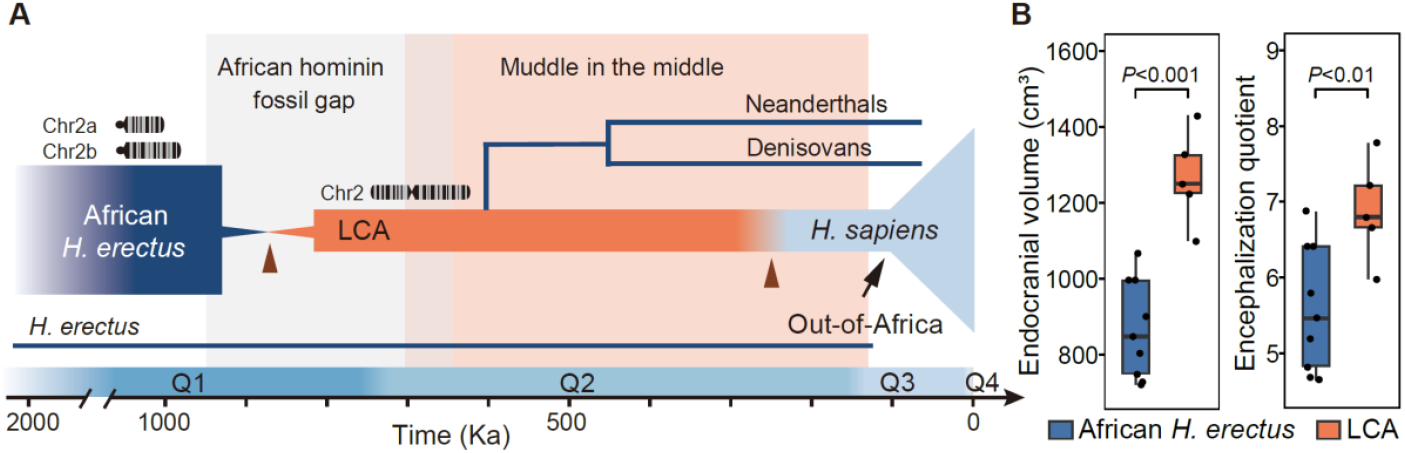
Integration of genetic evidence and fossil record. (**A**) Evolutionary history of hominins over the past one million years. LCA refers to the last common ancestor shared by Denisovans, Neanderthals, and modern humans. Triangles indicate speciation events. Q1 – Q4 denote the Early Pleistocene, Middle Pleistocene, Late Pleistocene, and Holocene, respectively. (**B**) Endocranial volumes and the encephalization quotients of African fossil specimens before and after the severe bottleneck. Central lines of boxplots represent the medians and box limits denote the 25th and 75th quantiles. Statistical significance was determined by two-tailed Student’s *t*-test.

In conclusion, high-precision calculation of expected SFSs is critical for minimizing artifacts in demographic inference. Recent studies emphasize that these considerations cannot be overlooked (Hilgers, et al. 2025; Ishigohoka and Liedvogel 2025). Collectively, advanced demographic inference and extensive human genomic data promise deeper insights into hominin speciation, genetic impacts of severe population bottleneck, and the evolutionary forces shaping modern humans. Our findings enhance the understanding of critical evolutionary events within the genus *Homo* and highlight the importance of integrating genetic, paleontological, and environmental evidence.

## Materials and Methods

The parameters of ancient bottleneck estimated from African and non-African populations were obtained from our previously published results (see the “Severe bottleneck during the Early to Middle Pleistocene transition” section) (Hu, et al. 2023). The student *t*-test was used to examine whether there is a significant difference between African and non-African estimates in Fig. 1B.

To illustrate the rich demographic information embedded within the SFS, we simulated 200 SFS datasets using *msprime* (Baumdicker, et al. 2022) under a simple bottleneck model in Figure 2C of Hu et al. Mutation and recombination rates were set at 1.*2* × 10^−8^ and 0.96 × 10^−8^ per base per generation, respectively, across 200 sequences of 100 Mb each. The generation time was 24 years. Demographic inference was performed with FitCoal, specifying three-time intervals for analysis. 10 repeats were performed to maximize the likelihood.

Four models were employed to investigate the effects of ancient severe bottlenecks on present-day human populations. Specifically, bottleneck models I and III from Hu *et al*. (Figure 4a and 4c) were utilized, with bottleneck I representing demographic history analogous to the true population history of African populations and bottleneck III analogous to that of non-African populations. Two additional models, modified versions excluding severe bottlenecks, were also derived from these original models. FitCoal calculated expected SFSs for all four scenarios, assuming *n* = 200 sequences.

We employed bottleneck model I as the true model, corresponding to the “severe bottleneck model” described by Deng *et al*. Using this model, we simulated 1,000 SFS datasets with *msprime*. Numbers of simulated sequences are 188, and the lengths of simulated sequences are 100 Mb each. These simulated datasets were then analyzed using *mushi* (DeWitt, et al. 2021) with parameters specified by Deng *et al*., yielding 1,000 inferred population histories. FitCoal was also used to infer population histories from the same simulated SFS datasets. Following our previous study, 10 repeats were performed to maximize the likelihood and analyze the simulated SFSs. Because FitCoal and *mushi* calculate the expected SFSs in different ways, to compare their likelihoods, we calculated the expected branch lengths for *mushi*-inferred demography using FitCoal.

The synthetic demographic model was obtained using the scripts from Deng et al. (Deng, et al. 2025). Initially, this model was generated using an expected SFS calculated by *mushi* based on a severe bottleneck scenario for an African population. The expected SFS was derived by multiplying branch lengths by the total mutation rate across the analyzed sequence fragments. To assess this synthetic model, we compared expected branch lengths for each SFS category using three independent methods. We assumed that *n* = 216 sequences. First, we calculated the expected branch lengths using *mushi*, following the approach described in recent studies (Cousins and Durvasula 2025; Deng, et al. 2025). Second, expected branch lengths were calculated using FitCoal version 1.3, freely available at https://doi.org/10.5281/zenodo.4765446 and released alongside this manuscript. Third, we performed simulations with *msprime*, generating 1 × 10^10^ coalescent trees for a single locus based on the synthetic model and calculating mean branch length per SFS category. The number of simulations 1 × 10^10^ was selected after evaluating expected branch length accuracy (Fig. 2E). Due to rounding precision considerations, at least 1 × 10^10^ coalescent simulations were necessary for accurate inference of expected branch lengths. These simulations were conducted on a CPU server equipped with an Intel Xeon Gold 6248R processor utilizing 60 parallel threads, with total runtime reported in core-hours. We loaded a pre-calculated table merely once to FitCoal, and FitCoal’s runtime for calculating expected branch lengths was determined as the average of 5,000 repetitions on the same server, using a single thread.

To examine the robustness of FitCoal in inferring population demography, we simulated 1,000 SFS datasets under the synthetic model using *msprime*. Simulation parameters matched those used by Deng et al. (2025), setting mutation and recombination rates at 1.*2* × 10^−8^ per base per generation across 216 sequences, each 100 Mb in length. Because the simulated data followed a neutral single-population model, exclusion of high-frequency mutations was unnecessary. FitCoal was run with default parameters, using a logLPRate of 20 (indicating a 20% likelihood gain) to infer 1,000 population size histories. Additionally, population history was inferred using a FitCoal-generated expected SFS with a logLPRate of 1 (indicating a 1% likelihood gain). The logLPRate setting determines the optimal model identification; a smaller logLPRate is appropriate when the expected SFS is used because maximum likelihood corresponds directly to the true model. However, since expected SFSs cannot be derived directly from samples and typically serve to calculate likelihoods for observed SFSs under demographic models, we recommend against using expected SFSs as input of FitCoal.

The endocranial volume data for the Buia specimen were sourced from Bruner et al. (2016), KNM-ER 42700 from Neubauer et al. (2018), and all remaining specimens from Holloway, Broadfield and Yuan (2004). Encephalization quotients for each specimen were calculated following Jerision (1973). Body mass estimates (in kilograms) used in these calculations were obtained from Carretero, et al. (2004), Grabowski, et al. (2015) and Jungers et al. (2016).

## Data Availability

FitCoal version 1.3 is freely available at https://doi.org/10.5281/zenodo.4765446.

## Acknowledgments

This work was supported by grants of National Natural Science Foundation of China to YHP (no. 31100273) and to HL (no. 32270674), the National Key Research and Development Project (2022YFF1203202, to HL), Eastern Talent Plan Leading Project (to HL), and CAS Youth Interdisciplinary Team (to ZS).

## Notes

### Competing Interest Statement

The authors have declared no competing interest.

### Summary of Updates

We have provided a new Table 1 to demonstrate the high accuracy of FitCoal. To further distinguish our work, we have also updated the title to "Severe bottleneck of ancient Homo populations".

